# Mindfulness meditation alters neural activity underpinning working memory during tactile distraction

**DOI:** 10.1101/790584

**Authors:** Michael Yufeng Wang, Gabrielle Freedman, Kavya Raj, Bernadette Mary Fitzgibbon, Caley Sullivan, Wei-Lin Tan, Nicholas Van Dam, Paul B Fitzgerald, Neil W Bailey

**Affiliations:** Monash Alfred Psychiatry Research Centre, Monash University Central Clinical School, Commercial Rd, Melbourne, Victoria, Australia; Epworth Centre for Innovation in Mental Health, the Epworth Clinic, Camberwell, Victoria, Australia, 3004; Brain and Mental Health Research Hub, School of Psychological Sciences, Monash Institute of Cognitive and Clinical Neurosciences, and Monash Biomedical Imaging, Monash University, Clayton, 3168 VIC, Australia; School of Psychological Sciences, The University of Melbourne, Parkville, VIC, Australia; Department of Psychiatry, Icahn School of Medicine at Mount Sinai, New York, NY, USA

**Keywords:** mindfulness, Alpha, P300, working memory, EEG

## Abstract

Evidence suggests that mindfulness meditation (MM) improves selective attention and reduces distractibility by enhancing top-down neural modulation. Altered P300 and alpha neural activity from MM have been identified and may reflect the neural changes that underpin these improvements. Given the proposed role of alpha activity in supressing processing of task-irrelevant information, it is theorised that altered alpha activity may underlie increased availability of neural resources in meditators. The present study investigated attentional function in meditators using a cross-modal study design, examining the P300 during working memory (WM) and alpha activity during concurrent distracting tactile stimuli. Thirty-three meditators and 27 healthy controls participated in the study. Meditators showed a more frontal distribution of P300 neural activity following WM stimuli (p = 0.005, η² = 0.060) and more modulation of alpha activity at parietal-occipital regions between single (tactile stimulation only) and dual task demands (tactile stimulation plus WM task) (p < 0.001, η² = 0.065). Additionally, meditators performed more accurately than controls (p = 0.038, η² = 0.067). The altered distribution of neural activity concurrent with improved WM performance suggests greater attentional resources dedicated to task related functions such as WM in meditators. Thus, meditation-related neural changes are likely multi-faceted involving both altered distribution and also amplitudes of brain activity, enhancing attentional processes depending on task requirements.

---

It is theorised that the effects of mindfulness-meditation (MM) on cognitive performance are mediated by improved neural resource allocation during stimuli processing, thereby prompting rapid attention allocation and reallocation speed, enhancing cognitive efficiency (Malinowski, 2013; Moore et al.,2012; van Leeuwen, Singer, & Melloni, 2012). Enhanced attentional processing, awareness, and sustained focus are considered to be core mechanistic components of many mindfulness-based practices (Lippelt, Hommel, & Colzato, 2014; Lutz, et al., 2008), and MM has been demonstrated to improve selective attention and reduce distractibility (Moore, Gruber, Derose, & Malinowski, 2012; Wong, Teng, Chee, Doshi, & Lim, 2018; Zeidan, Johnson, DiamondDavid, & Goolkasian, 2010). These changes result in enhanced performance of resource demanding cognitive processes, evidence of which has been shown in tasks requiring executive functions, memory, and self-regulation (Chiesa, Calati, & Serretti, 2011; Manna et al 2010; Tang & Posner, 2009). These improvements may be related to altered neural activity (Fox et al., 2016) and structure (Fox et al., 2014) in meditators and provide insights into the underlying neural basis of attentional changes through MM. An important neural marker of mindfulness related performance enhancements has been found in electroencephalography (EEG) studies using event-related-potentials (ERPs) exploring task-dependent P300 modulation (Slagter et al., 2007). The P300 is a positive voltage deflection typically peaking around 300ms after presentation of a target stimulus. It is likely related to activity in the frontal and temporal-parietal networks and functions to facilitate selective attention and visual working memory (WM; Polich, 2007). The P300 is considered an index of resource allocation during complex task performances and its amplitudes have shown correlations with increased demand for evaluative resources (Kok, 2001; Polich, 2007; Slagter et al., 2007). Meditators have shown reductions in attentional-blink effects (where the second stimulus in a sequence of rapidly presented stimuli is commonly missed; Slagter et al., 2007), along with reductions in P300 amplitudes to the first stimuli (Slagter et al., 2007). Mindfulness practice has also been associated with reduced interference from distractor stimuli and greater control of resource allocation across multiple modalities (auditory and visual; van den Hurk, Giommi, Gielen, Speckens, & Barendregt, 2010), more efficient neural processing and attentional network functioning (Isbel, Lagopoulos, Hermens, & Summers, 2019), and improved ability to direct attentional resources toward task relevant stimuli (Moore et al., 2012).

Although evidence suggests mindfulness practice can influence neural resource allocation, the mechanism by which this occurs is unclear. One theory proposes that mindfulness related attention enhancements are the result of practice-specific effects on alpha modulation (Kerr, Sacchet, Lazar, Moore, & Jones, 2013). Modulation of the alpha rhythm has been observed to regulate sensory inputs to the somatosensory cortex and is seen as a filtering mechanism in a range of information processing tasks (Foxe and Snyder, 2011). For example, alpha modulation has been found to correlate with WM load and performance, with greater alpha power over sensory processing areas during higher memory load, and greater alpha power associated with higher probabilities of correct WM responses (Jensen, Bonnefond, & VanRullen, 2012; Jensen and Mazaheri, 2010; Scheeringa et al., 2009). Additionally, modulation of alpha in the primary somatosensory cortex appears to facilitate sensory throughput when people are cued to direct attention to specific body regions (Pritchett, Jones, Wan, Moore, Kerr, & Hamalainen, 2010). Thus, modulation of alpha over sensory regions appears to act as a suppression mechanism during resource demanding cognitive processes by reducing distractor processing (Sauseng et al., 2009). Mindfulness-meditation is likely to facilitate more efficient cognitive performance during resource demanding tasks (such as WM) by supressing irrelevant stimuli processing (Kerr et al., 2013), and decreased processing of task-irrelevant information among meditators is likely related to increased alpha modulation.

At the present, preliminary evidence suggests that mindfulness related cognitive enhancements may be enabled by 1) increased attention to the target stimuli (with increases in associated neural activity), 2) increased suppression of task-irrelevant sensory information, potentially via higher alpha activity over task-irrelevant cortical regions, or 3) a combination of both. However, previous methods have not permitted dissociation of these possibilities. A better understanding of the mechanism by which mindfulness exerts its effects could facilitate a better understanding of when and for what conditions it might be helpful.

The current study used a cross-modal design combining a WM task with a tactile distractor to allow for specific testing of the role of target-enhancing and distractor-suppressing mechanisms in attentional processing. The study investigated differential processing of concurrently presented visual and somatosensory stimuli in meditators and controls by examining P300 ERPs related to visual WM stimuli and alpha activity related to tactile stimuli. In particular, we tested whether meditators would demonstrate enhanced WM related electrophysiological responses concurrent with greater ability to suppress sensory processing (reflected by higher alpha activity over somatosensory regions). We predicted that meditators would show enhanced ERP amplitudes to WM stimuli, reflecting enhanced attention despite distraction, irrespective of whether distractors required a response. Secondly, research has indicated that meditators demonstrate more pronounced frontal ERPs when attending to task-relevant stimuli, reflecting more engagement of attention-relevant areas (Bailey et al. 2018). As such, we expected WM-related ERPs in the current study to show more frontal distributions among meditators (again irrespective of whether distractors required a response). Thirdly, preliminary evidence has indicated enhanced ability in meditators to modulate alpha activity in regions processing distractor stimuli (Kerr et al., 2013). With the current study design, this effect was tested in two ways. First, when only tactile stimuli were presented (no visual stimuli), increased alpha activity was expected in visual processing regions in meditators, reflecting suppression of non-relevant visual processing regions. Secondly, when WM stimuli were presented concurrent with a tactile distractor, increased alpha activity in somatosensory processing regions was expected in meditators, reflecting suppression of non-relevant somatosensory information.

Our primary hypotheses were as follows. 1) Meditators were expected to show larger P300 amplitudes toward visual WM stimuli than the control group, indicating increased processing of WM stimuli. 2) Meditators were expected to show a more frontal distribution of the P300 to WM stimuli reflecting increased engagement of attentional processing regions. 3) Meditators were expected to show greater alpha activity over somatosensory regions (time locked to tactile stimuli) during the visual WM task than controls. 4) Meditators were expected to show greater alpha activity over visual processing regions during task conditions requiring tactile-only processing, reflecting suppression of task irrelevant brain regions for additional neural resources. Exploratory source analyses was planned to characterise the source of neural activation differences between groups (without statistical comparisons) and exploratory microstate analyses were planned to characterise the shifting pattern of neural activity across the period after WM stimuli presentation. Lastly, as neural data was the focus of this study, behavioural comparisons were exploratory without specific directional hypotheses.

## Methods

### Participants

Seventy participants between the ages of 18 to 65 (45 females and 25 males) were recruited for the study; 34 mindfulness meditators and 36 healthy control non-meditators. Participants were recruited via community, meditation centres, and university advertising, and were reimbursed a total $30 for their participation.

Meditation participants were included if they had practiced for more than two years and currently practiced more than two hours per week. Screening was conducted through phone and in-person interview by experienced mindfulness researchers (GF, KR, NWB), to ensure mindfulness practice were congruent with Kabat-Zinn’s definition - “paying attention in a particular way: on purpose, in the present moment, and nonjudgmentally” (Kabat-Zinn, 1994). Screening also ensured meditation practices were consistent with either focused attention on the breath or body-scan. Uncertainties were resolved through discussion between the principle researcher (NWB) and a second researcher. Control group participants were excluded if they reported more than two hours of lifetime meditation experience.

Exclusion criteria included self-reported current or past experiences of mental or neurological illness, current psychoactive medication or recreational drug use. These were further assessed using the Mini International Neuropsychiatric Interview for DSM-IV (Hergueta, Baker, & Dunbar, 1998), Beck Depression Inventory-II (BDI-II), and Beck Anxiety Inventory (BAI) (Beck & Clark, 1997; Beck, Steer, & Brown, 1996) which were administered by GF, KR, or NWB. Participants were excluded if they met diagnostic criteria for any DSM-IV psychiatric disorders, or if they scored in the mild or above range on the Beck anxiety and depression scales.

Prior to completing the task, participants provided demographic information and reported their estimated years of mindfulness practice and minutes per week of current practice. Self-report measures were also completed; Freiburg Mindfulness Inventory (FMI) (Walach et al., 2006), Five Facet Mindfulness Questionnaire (FFMQ) (Baer et al. 2006), BAI, and BDI. Written informed consent was obtained from participants prior to the commencement of the study. The Monash University Human Research Ethics Committee and the Alfred Hospital Ethics Committee approved all experimental procedures.

Four participants in the control group were excluded due to high scores on the BDI (within clinical range) and one control was excluded due to task non-completion. In order to maximise data available for analysis, further exclusions of select data were made for neural analysis separately. Four controls and one meditator were excluded from the neural analysis, as they provided too few artefact free EEG epochs for analysis (based on the criteria provided in the Electrophysiological Recording and Data Processing section). Thus, final analyses were run on 27 controls and 33 meditators for neural analysis and 31 controls and 34 meditators for behavioural analysis.

### Procedure

To study the effect of cross-modal task demands on attention and related neural activity in meditators, we used a 2 group (meditators vs controls) × 2 condition (attend/ignore tactile stimulation) design across 2 task conditions with different sensory modalities (visual working memory/tactile oddball), with four within subject conditions in total (Figure 1). Each participant was tested in one continuous session split in four different testing conditions while EEG was recorded (details of the EEG recording below). In the first condition, participants experienced brief tactile sensations (comprising a tactile oddball), which they were instructed to ignore (condition [C] 1: Ignore Tactile Only). In the second condition, participants continued to experience and were instructed to ignore the tactile sensations, while concurrently performing an N-back WM task (2-back), pressing button 1 for target letters (Condition 2: Ignore Tactile, Attend N-back). In the third condition, participants experienced tactile sensations without the N-back task being present, and were instructed to respond by pressing button 2 after the infrequent occurrence of two sensations presented in close temporal proximity (condition 3: Attend Tactile Only). The final condition involved participants responding with one button to the N-back task, and with another button to the infrequently presented double tactile stimuli (condition 4: Attend Tactile, Attend N-back).

**Figure 1.**
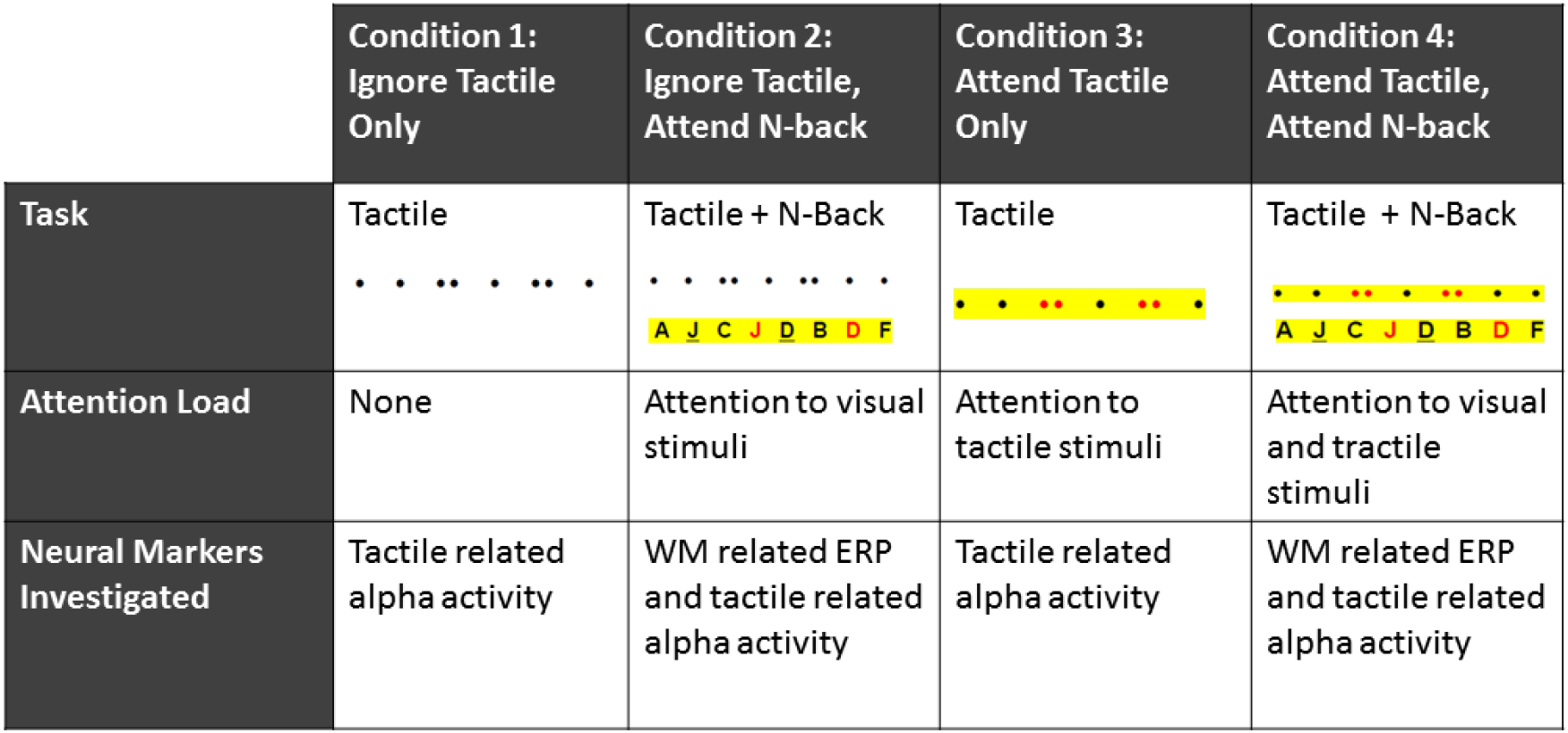
Task design. All participants completed conditions 1-4 in the same order. Dots represent the tactile stimulation; single dots represent a single tactile stimulation, double dots represent a target oddball double tactile stimulation, red double dots highlight the target during attend conditions (requiring response with button 2). Letters represent the N-back WM letters; underlined letters represent the first letter in the target pair, red letters represent memory targets to which a response is required (with button 1). Yellow represents the direction of attention according the task instructions.

In the first testing condition, participants were administered 125 brief electrotactile stimuli and instructed to remain quietly seated while ignoring the stimuli by directing their gaze toward the screen in front without any particular attentional focus (condition 1: Ignore Tactile Only). The electrotactile stimuli were administered to the anterior radial portion of the left wrist (median nerve) using a transducer with two flat metal probes (cathode proximal) of 9mm diameter at 30 mm spacing apart (Stimulating Bar Electrode, MLADDF30, AD Instruments, NSW, Australia). Electrical current was passed between these two probes for 80ms during each pulse and was delivered by digitally generated waveforms converted to analogue signal and then amplified (Powerlab, AD Instruments, FE116, NSW, Australia). The amplitude of electrical stimulation was set at 1.75x the sensory threshold (which was determined by delivering randomly timed brief electrical stimulations at amplitudes slowly increasing from 0.5 volts until participants consistently reported detecting the sensation, then reducing the amplitude until participants could no longer detect the sensation). The sensory threshold was defined as the amplitude 0.1 volts above the value at which participants reported detecting the sensation with 100% accuracy. Amplitude was set at 1.75x the sensory threshold stimulation value to account for variation in participants’ sensory threshold. Participants confirmed the sensation as noticeable but not eliciting discomfort, and all participants included in analyses provided responses to the tactile stimuli in conditions where they were required to do so, confirming that the sensations were detected. Amplitude of the electrical signal varied between 0.7 and 2.3 volts depending on each individual’s threshold. Oddball stimuli consisted of double stimulations (with an inter-stimulus-interval of 250ms) and were randomly dispersed throughout the series accounting for 16% of the trials. The same stimulation was used to deliver both single standard stimulations (random inter-trial interval of 1800-2840ms, with an average of 2320ms) and oddball stimulations.

During the second condition, the N-back WM task was introduced, and participants were asked to attend to the N-back task while continuing to ignore the tactile stimulation (the 125 regular and oddball tactile stimulations were presented through the second condition; Condition 2: Ignore Tactile, Attend N-back). The N-back memory task was presented on a computer approximately 80cm in front of participants in a darkened and sound attenuated room. A series of letters (from A to J) were presented in random order. Participants responded by pressing button 1 when the currently viewed letter was the same as the letter presented two trials previously (2-Back). A short practice task was provided prior to initiation of the real task. The complete N-back task consisted of 390 visual WM stimuli, in two blocks of 195, with a 1.5 second inter-trial interval and trials contained 25% of target letters. The second block was designed to provide a measure of neural activity related to WM during ongoing tactile distraction (described below). The timing of tactile stimulations had no relationship to the presentation of memory stimuli in all blocks and was identical to the sequence in the Ignore Tactile Only condition in all conditions.

The third condition presented the tactile oddball, but not the N-back task. Participants were required to attend to the 125 tactile stimuli and press button 2 when they felt the oddball double stimulation (condition 3: Attend Tactile Only). The fourth block presented both the 125 tactile stimuli and the N-back stimuli concurrently. Participants were asked to continue attending to the tactile stimuli pressing 2 for an oddball double stimulation, and at the same time perform the N-back task, pressing button 1 for a target letter (condition 4: Attend Tactile, Attend N-back). In this fourth condition, participants were asked to split their attention between dual tasks, allowing measurement of WM ERPs and alpha activity to tactile stimulation during an attention-limited state with greater task demands. Task transitions were demarcated by new instructions informing participants of changes in the task requirements.

### Data Analysis

#### Behavioural Data Comparisons

Tactile oddball and N-back response accuracies were compared by first calculating d-prime scores (*d*’=*z*[hit rate]-*z* [false alarm rate]) as this measure allows for more accurate comparison of differences in behavioural performance (Wickens, 2002). *d*’ scores were then analysed for each type of stimuli separately using a repeated measures ANOVA in SPSS, as were reaction times to both types of stimuli separately. *d*’ was compared using a repeated measures ANOVA for response to N-back letters (2 groups x 2 conditions – C2: Ignore Tactile, Attend N-back/C4: Attend Tactile, Attend N-back)) and for response to tactile stimulation (2 groups x 2 conditions – Attend C3: Tactile Only/C 4: Attend Tactile, Attend N-back). Reaction time was compared using a repeated measures ANOVA for response to N-back letters (2 groups x 2 conditions – C2: Ignore Tactile, Attend N-back/C4: Attend Tactile, Attend N-back) and for response to tactile stimulation (2 groups x 2 conditions – C1: Attend Tactile Only/C4: Attend Tactile, Attend N-back). No outliers were detected (more than 3.29 SD from the mean). There was no evidence for violation of univariate or multivariate equality of variance nor for violations of normality.

#### Electrophysiological Recording and Data Processing

A 64-channel Neuroscan EEG Ag/AgCl Quick Cap acquired data to Neuroscan software through a SynAmps2 amplifier (Compumedics, Melbourne, Australia). Electrodes were referenced online to an electrode between Cz and CPz. Horizontal and vertical eye movements were recorded using four EOG electrodes located above and below the left orbit and adjacent to the outer canthus of each eye. Impedances were maintained at less than 5kΩ. Recordings were sampled at 1000 Hz and bandpass filtered from 0.05 to 200Hz (24 dB/octave roll off). MATLAB (The Mathworks, Natick, MA, 2016a) and EEGLAB were used for pre-processing of EEG data (sccn.ucsd.edu/eeglab; Delorme and Makeig, 2004). Second-order butterworth filtering was applied to the data with a bandpass from 1-80Hz and a band stop filter 47-53Hz. Data was then epoch time-locked to the onset of the single pulse tactile stimulation (-1000 to 2000ms) and also epoch time-locked to the onset of the visual stimuli in the WM conditions (-1000 to 2000ms).

All analyses of alpha activity were performed on data time-locked to single pulse tactile stimuli rather than to oddball stimuli. The task design presented single pulse stimuli at greater frequency than the oddball stimulus, this provided more neural data and allowed more reliable interpretations. Participants were not instructed to respond to the single pulse tactile stimuli, and epochs containing responses to single pulse tactile stimuli or to the N-back stimuli were excluded to avoid confounding motor activity. Epochs time-locked to the onset of visual WM stimuli also excluded epochs with responses. Epochs were visually inspected by an experimenter experienced with EEG analysis and periods containing muscle artefact or excessive noise were excluded as were channels with low quality signals. Each participant provided 35 or more accepted epochs for each condition, and no significant differences were detected in the number of accepted epochs for each block (all p’s > 0.10). Adaptive mixture independent component analysis was used to manually select and remove components related to eye movement and remaining muscle activity (Palmer, Kreutz-Delgado, & Makeig, 2011). Once removed, data was then re-filtered from 0.1 to 80 Hz and was again inspected by a separate researcher blinded to the group identity of data inspected at the time. Recordings were re-referenced offline to an averaged reference. Epochs were then averaged within each WM condition and each participant for statistical analysis of ERPs. Alpha activity related to tactile stimuli was computed using a Mortlet Wavelet multi-taper convolution transform with a 3.5 cycle width and a Hanning taper to provide a measure of power in the 8-13 Hz alpha frequency range. These power values were averaged across epochs for each of the conditions separately for each participant.

#### Statistical Comparisons

Self-report and behavioural results were analysed using SPSS version 23. Independent samples t-tests were performed to examine potential group differences in age, years of education, BDI, and BAI, FMI and FFMQ scores. Potential differences in categorical data (gender and handedness) were examined using the Chi square test.

#### Primary Comparisons

Statistical comparisons of EEG data were conducted using the Randomization Graphical User Interface (RAGU), which uses rank order randomization statistics to compare scalp field differences from all electrodes and epoch time points between groups (Koenig, Kottlow, Stein, & Melie-García, 2011). The software uses the spatial standard deviation of the electric field from all electrodes simultaneously to obtain a single Global Field Power (GFP) value for each point in time across the epoch, thus controlling for multiple comparisons across space. GFP is a reference-free EEG measure, which avoids an arbitrary choice of reference (Koenig et al., 2011). Ragu also allows for the comparison of scalp distributions of neural activity with the Topographic Analysis of Variance (TANOVA). The recommended L2 normalization was performed to normalize differences in individual neural response amplitude, so that TANOVA comparisons test differences in the distribution of neural activity independently of differences in amplitude.

Additionally, the topographical Consistency Test (TCT) compared GFPs of each participant to randomly shuffled data to assess whether there is consistent topographical activation within groups (Koenig and Melie-García, 2010). Inconsistent topographical activation can be interpreted as within group variability and supports the null hypothesis of no difference from 0 signal within a group/condition, whereas consistent topographical activation within groups allows valid comparisons between groups and conditions (Koenig and Melie-García, 2010).

Comparisons of ERP data related to WM stimuli were made for the entire 0 to 1000ms window following the stimuli. GFP and TANOVA tests were used to conduct 2 group × 2 condition (C2: Ignore Tactile, Attend N-back/C: Attend Tactile, Attend N-back) comparisons for WM ERPs. Comparisons of alpha power were made using averaged activity over the 0 to 1000ms window following the onset of the tactile stimulus. Regarding alpha comparisons, it should be noted that when frequency transformed data comparisons are performed with RAGU, the average reference is not computed with the transformed data (the average reference was computed prior to the transforms). As such, the test is a comparison of the Root Mean Square (RMS) between groups, a measure that is a valid indicator of neural response strength in the frequency domain. In other respects, the statistic used to compare RMS between groups is identical to the GFP test described in the previous paragraph. Alpha values were compared with Root Mean Square (RMS) and TANOVA tests (to separately compare overall neural response strength and distribution of neural activity, respectively). RMS and TANOVA tests were used to conduct 2 group x 2 condition (C2: Ignore Tactile, Attend N-back/C4: Attend Tactile, Attend N-back) x 2 condition (C1: Ignore Tactile Only/C3: Attend Tactile Only) comparison for tactile related alpha activity.

The recommended 5000 randomization runs were employed for each statistical test. Global duration statistics were used to control for multiple comparisons across time for the ERP data (alpha comparisons did not require multiple comparison control in the temporal dimension as alpha activity was averaged across the 0 to 1000ms window after stimuli presentation). Global duration statistics calculated the duration of significant effects within the epoch that are longer than 95% of significant periods in the randomized data. This ensures that significant differences in the real data last longer than the random comparison data with our alpha level of 0.05 (Grieder et al., 2012). Refer to Koenig and Melie-Garcia (2010) and Koenig et al. (2011) for further information regarding these analyses.

The Benjamini and Hochberg false discovery rate (FDR) (Benjamini and Hochberg, 1995) was performed on the global count p-values from each main effect or interaction. The FDR reduces the false discovery rate and is used to control multiple comparisons for all comparisons involving the primary hypothesis separately from behavioural comparisons. P values are labelled ‘FDR p’ and ‘p-uncorrected’ to allow for comparison with other research.

#### Exploratory Analysis

Source analysis and microstates analysis were used to further explore differences in ERPs following visual stimuli. Microstates are transient patterns of scalp topography lasting from milliseconds to seconds before transitioning to another temporarily stable topography and are hypothesised to be the basic building blocks of neural functioning (Koenig et al., 2002). Ragu was used to identify and analyse microstates during significant windows. Results of the source and microstate analysis are reported in the supplementary materials.

## Results

### Demographics

Neural analyses were the focus of this study, so only participants selected for neural analyses were examined for differences in demographic and self-report data. Results are summarised in table 1. No significant differences were found between groups in age, BAI and BDI scores, gender or handedness (all *p* > 0.3). However, meditators scored significantly higher on the FMI, *t*(58) = 2.401, p = 0.019, and FFMQ, *t*(58) = 3.741, p < 0.001, compared to controls. Meditators also exhibited more years of education *t*(58) = 2.01, *p* = 0.049. As the difference between groups could be a confound, we replicated all significant comparisons after excluding the six meditators with the highest number of years of education. After exclusion of those six meditators, groups did not differ in years of education (*t*(52) = 0.925, *p* = 0.359). All significant results remained significant when groups without differences in years of education were compared (*p* < 0.05, with larger effect sizes found in all tests, reported in supplementary materials).

**Table 1.** Demographics and self-report data.

|  | Meditators *M(SD)* | Control *M(SD)* | Statistics |
| --- | --- | --- | --- |
| N | 33 | 27 |  |
| Sex(F/M) | 21/12 | 17/10 | $\chi^2(1) = 0.0029, p = 0.957$ |
| Age | 36.91 (10.85) | 35.07 (13.87) | $t(58) = 0.575, p = 0.568$ |
| Years of Education | 17.09 (2.49) | 15.79 (2.45) | $t(58) = 2.011, p = 0.049^*$ |
| Meditation Experience<br>(years) | 8.43 (10.41) | 0 |  |
| Current Meditation Practice<br>Per Week (hours) | 5.24 (3.93) | 0 |  |
| BDI score | 1.09 (1.89) | 1.67 (2.69) | $t(58) = 0.971, p = 0.336$ |
| BAI score | 4.33 (4.72) | 4.78 (5.55) | $t(58) = 0.335, p = 0.739$ |
| FMI score | 45.75 (7.08) | 40.93 (8.06) | $t(58) = 2.470, p = 0.016^*$ |
| FFMQ score | 153.58 (17.02) | 138.11 (12.83) | $t(58) = 3.899, p < 0.001^{**}$ |

\* $p < 0.05$ , \*\* $p < 0.001$ .

### Behavioural Performance

For N-back results, a significant main effect of group was found for *d*’, with meditators performing more accurately than controls (F(1,63) = 4.516, *p* = 0.038, η² = 0.067). No significant main effect was found for reaction time nor for the interaction between Group x visual WM conditions (C2: Ignore Tactile, Attend N-back/C4: Attend Tactile, Attend N-back conditions for reaction time or *d*’ (all *p* > 0.5). Additionally, for responses to the tactile stimulation, no main effect of group nor interaction between Group x Tactile conditions (C1: Attend Tactile Only/C4: Attend Tactile, Attend N-back conditions) were found in *d*’ or reaction time (all *p* > 0.07). Figure 2 presents d’ scores and reaction times in behavioural performances. For a complete table of means, standard deviations and statistics for behavioural performance comparisons please see Supplementary Table 1.

**Figure 2.**
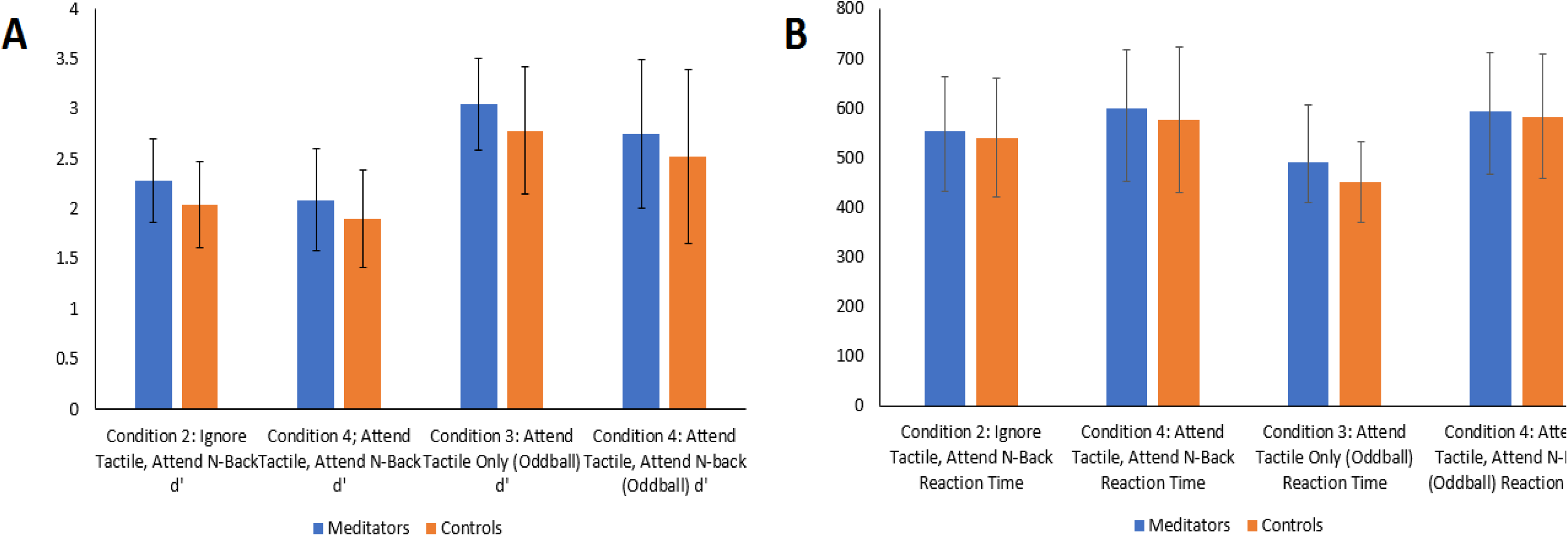
Signal detection (*d’*) scores and reaction times in behavioural performance. Panel A. Mean N-Back and tactile oddball *d’* scores. Error bars represent standard deviation. Panel B. Mean N-back and tactile oddball reaction times. Error bard represent standard deviation

### Neural Data; Visual Stimuli Locked ERPs

#### TCT and GFP

The TCT (Koenig et al., 2011) showed topographical consistency within all groups and conditions indicating that TANOVA comparisons are valid during the majority of time periods (see supplementary materials 1 for a more detailed description). The GFP randomization test was performed to assess the strength of ERP neural response to N-back stimuli in C2: Ignore Tactile, Attend N-back and C4: Attend Tactile, Attend N-back conditions. There was no significant main effect of Group (global count statistics across the whole epoch *p* = 0.280, FDR *p* = 0.350), nor interaction for Group x C2: Ignore Tactile, Attend N-back/C4: Attend Tactile, Attend N-back conditions in GFP (no periods of significance lasted longer than the duration control of 47ms, global count statistics across the whole epoch *p* = 0.198, FDR *p* = 0.282). See figure 3.

**Figure 3.**
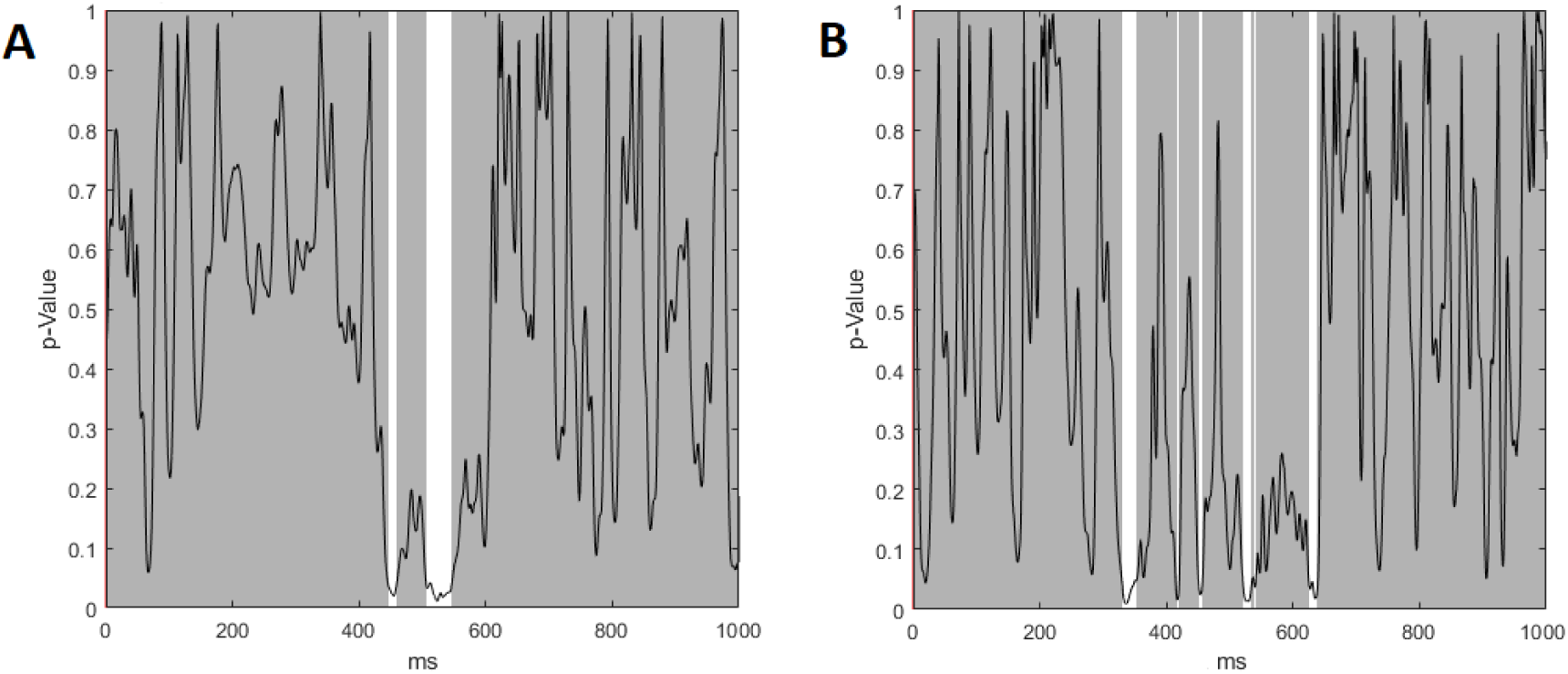
A- Main effect of group in GFP test across the duration of the epoch. B- Group effects by N-Back conditions (C2: Tactile, Attend N-back/C4: Attend Tactile, Attend N-back) interaction in GFP test across the duration of the epoch. No significant main effect of Group, nor interaction for Group x C2: Tactile, Attend N-back/C4: Attend Tactile, Attend N-back conditions in GFP were present (no significant periods lasted longer than the duration control of 47ms).

#### TANOVA

TANOVAs were conducted to examine neural activity distribution in response to the N-back stimuli during the visual WM conditions (C2: Ignore Tactile, Attend N-back/C4: Attend Tactile, Attend N-back conditions). The main effect of group comparison showed a significant difference during two separate time windows, with meditators showing a P300 with a more frontal distribution of positive voltages during both time windows (global count statistics across the whole epoch *p* = 0.009 FDR *p* = 0.045). A significant main effect of group that survived duration control (44ms) for multiple comparisons was found in TANOVA from 299 to 352ms. This result was significant when activity was averaged across a significant window from 299 to 352ms (*p* = 0.005, η² = 0.060).

A significant effect of group was also found in TANOVA from 419 to 613ms, however in light of the TCT results which showed within group variance among meditators (527-606ms in C4: Attend Tactile, Attend N-back condition, and from 524-580ms in C2: Ignore Tactile, Attend N-back condition), the main effect of group from 524ms onwards could be due to within group/condition variability. Thus, from 419 to 524ms there was a significant main effect between groups (*p* = 0.001, η² = 0.076, global duration control statistic was 44ms), but after this period, apparent differences between groups may be attributed to variance within the meditator group. Figure 4 depicts topographical differences between groups for the first significant window from 299 to 352ms, and for the second significant window from 419 to 524ms.

**Figure 4.**
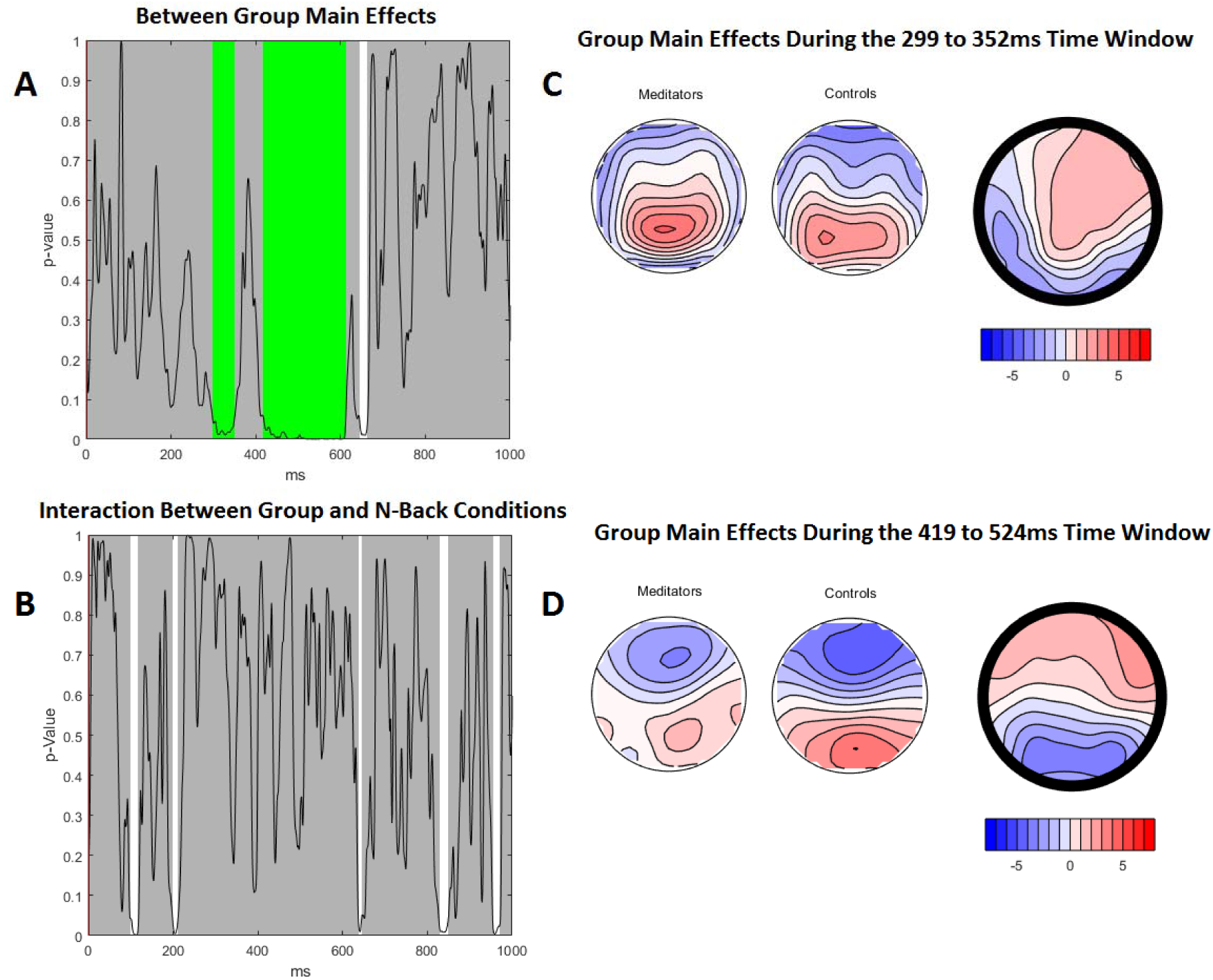
TANOVA main group effect. A– p-graph showing the significant main effect of group and p-values across the duration of the epoch (global count statistics across the whole epoch *p* = 0.009, FDR *p* = 0.045). The probability of the null hypothesis was below 0.05 from 299 to 352ms and from 419 to 613ms. Green bars reflect periods that exceeded the duration control for multiple comparisons across time (44ms). This period was a longer duration of significance than 95% of the 5000 randomizations. B– Group effects by visual WM conditions (C2: Ignore Tactile, Attend N-back/C4: Attend Tactile, Attend N-back) across the duration of the epoch. *p* > 0.05 for the entire period, except for a few brief time periods but did not last the duration control for multiple comparisons across time (44ms; count statistics across the whole epoch *p* = 0.11). C – topographic maps for each group and a *t*- map for meditators topography minus control topography during the 299 to 352ms time window (*p* = 0.005, averaged across the first significant period, η² = 0.060). D - topographic maps for each group and a t- map for meditators topography minus control topography during the 419 to 524ms time window (p = 0.001, averaged across the second significant period, η² = 0.076).

Overall, the differences indicate a P300 with a more central and right frontal-central positivity in the meditation group from 299 to 352ms, and positive voltages extended further frontally in the meditation group from 419 to 524ms. Because the P300 typically shows maximal positivity over parietal areas on the scalp and is thought to peak at around 300ms, the difference in this activity may reflect more frontal regions activated in the meditation group during performance of the function the P300 is associated with - working memory and context updating processes (Kok, 2001; Polich, 2007).

No interaction between group and visual WM conditions (C2: Ignore Tactile, Attend N-back/C4: Attend Tactile, Attend N-back) was present (p > 0.05 for the entire time period, except for a few brief time periods that did not last longer than duration control multiple comparisons, global count statistics across the whole epoch p = 0.106; FDR p = 0.266).

### Neural Data; Tactile stimulation locked alpha

#### Root Mean Square Test

The RMS test was performed to assess the strength of neural response within the alpha frequency averaged across the 0 to 1000ms period for all conditions. No significant main effect of group was present (*p* = 0.911, FDR *p* = 0.974), nor interaction between group and Condition 2/Condition 4 (ignore/attend tactile stimuli; *p* = 0.197, FDR *p* = 0.282). A significant interaction was found between group and visual WM N-back task absent/present conditions (C3: Attend Tactile Only/C4: Attend Tactile, Attend N-back) which did not survive FDR multiple comparison control (*p* = 0.017, FDR *p* = 0.057, η² = 0.037, ηp²= 0.062). For post-hoc analysis, N-back conditions were averaged together (C2: Ignore Tactile, Attend N-back/C4: Attend Tactile, Attend N-back) and tactile only conditions were averaged together (C1: Ignore Tactile Only and C3: Attend Tactile Only). Post-hoc analysis between groups for averaged N-back conditions (*p* = 0.3714,) and averaged tactile only conditions (*p* = 0.4854) were both not significant. However, significant differences were found between the N-back and tactile only conditions within both meditator (*p* = 0.0002, η² = 0.486) and control groups (*p* = 0.0002, η² = 0.277). It seems that the interaction is driven by the control group showing less difference between the two grouped conditions. RMS test results for the interaction between group and the WM task present/absent conditions are presented in figure 5.

**Figure 5.**
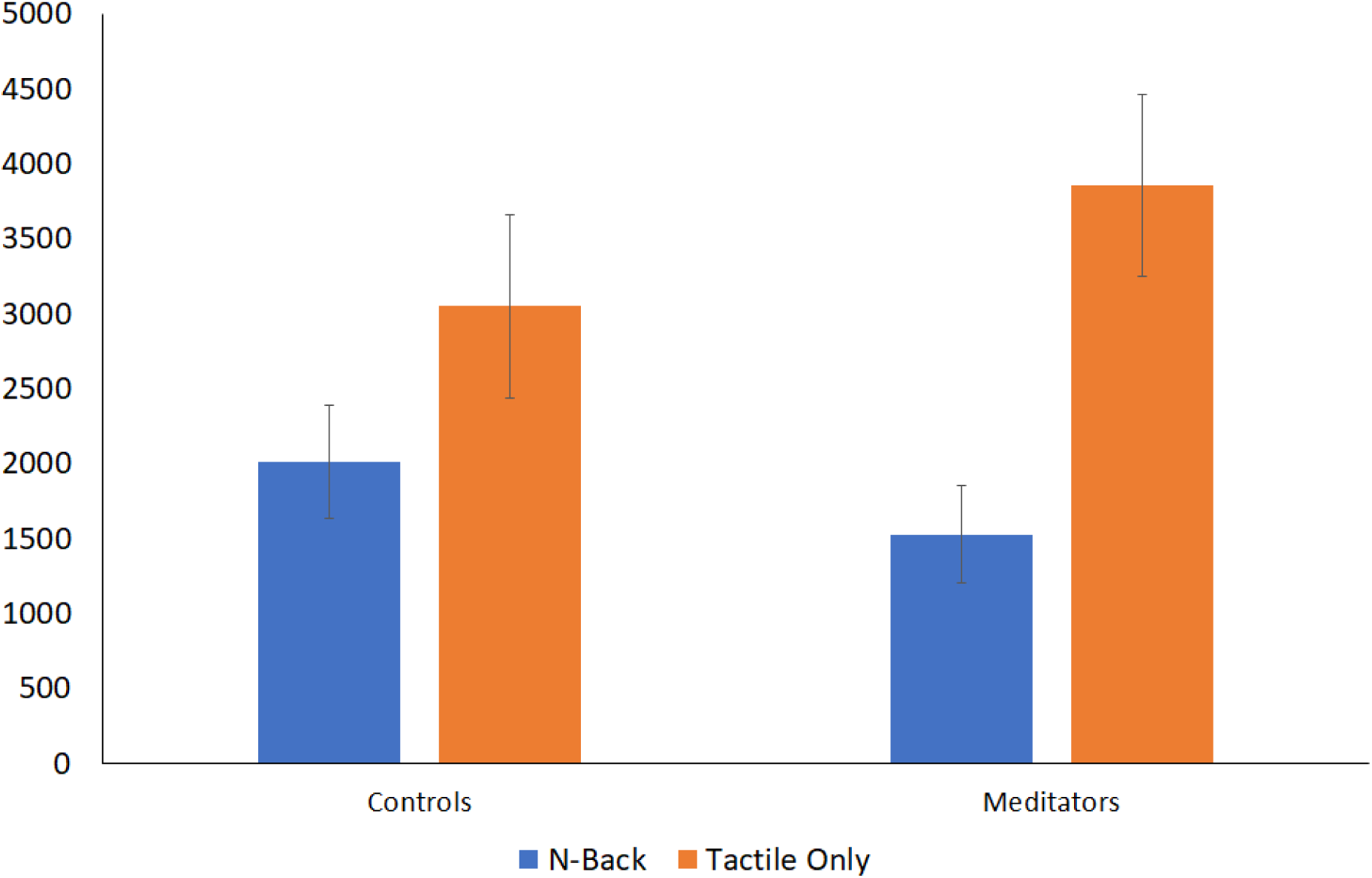
Averaged alpha activity RMS values across the significant time period 0 to 1000ms. Meditators showed a greater difference between conditions than controls (*p* = 0.017, FDR *p* = 0.057, η² = 0.037, η² = 0.062). Significantly larger RMS values in the averaged Tactile Oddball Only conditions than averaged N-back conditions were found for both meditator (*p* = 0.0002, η² = 0.486) and control groups (*p* = 0.0002, η² = 0.277). *Note*: N-Back = average of conditions with the N-back Task (Conditions 2 and 4), Tactile Only = average of conditions without the N-Back task (Conditions 1 and Condition 3).

#### TANOVA

No significant main effect of group was present (*p* = 0.154, FDR *p* = 0.329), nor main effect of tactile stimuli being ignored/attended (p = 0.010). Additionally, no significant interaction was found between group and whether tactile stimuli were ignored/attended (C2: Ignore Tactile, Attend N-back/C4: Attend Tactile, Attend N-back; *p* = 0.974, FDR *p* = 0.974). A significant interaction was found between group and whether participants were completing the N-back memory task or not (C3: Attend Tactile Only/C4: Attend Tactile, Attend N-back; *p*-uncorrected < 0.001, FDR *p* = 0.002, η² = 0.056, ηp² = 0.065). Post-hoc results suggested compared to controls, meditators showed an altered distribution of neural activity with more posterior alpha in the Tactile Only conditions (*p* = 0.047), with no significant difference for the N-back conditions (*p* = 0.155). Furthermore, meditators showed greater posterior alpha in the Tactile Only conditions compared to N-back conditions (*p* < 0.001, η² = 0.185; see figure 6). Although controls also differentiated significantly between conditions (*p* = 0.046, η² = 0.04), this change was not as strong as the differences found for meditators.

**Figure 6.**
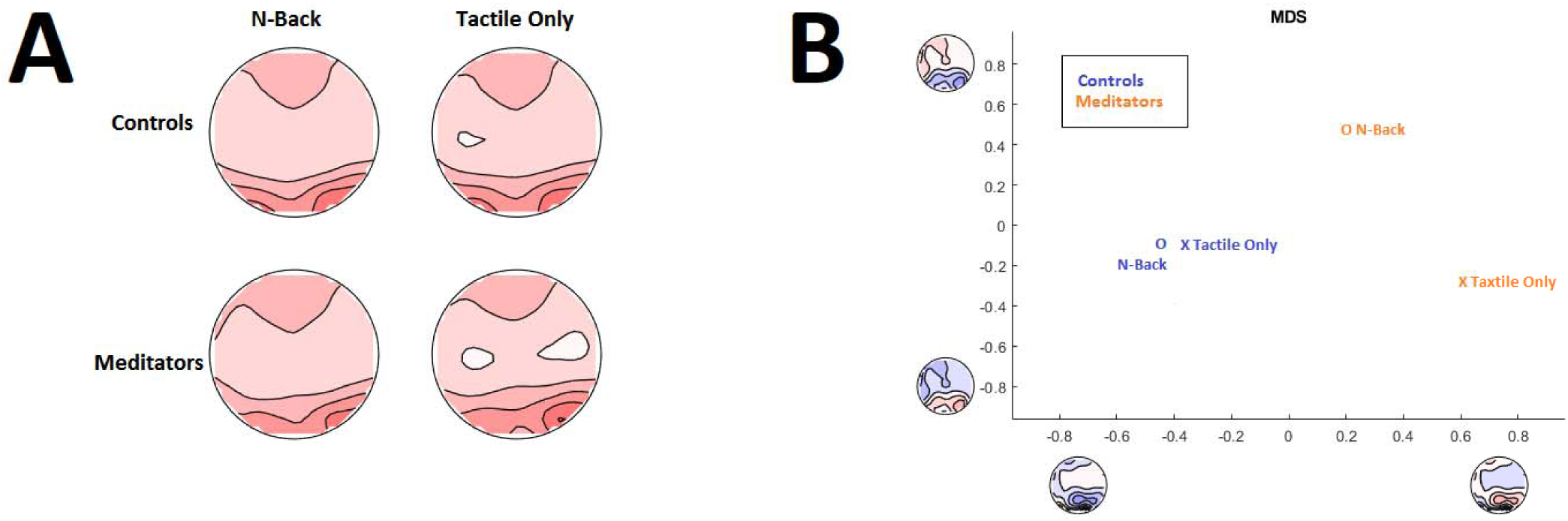
A - Averaged alpha topographies of Group x N-back/Attend Tactile Only conditions averaged across the 0 to 1000ms window. Meditators showed an altered distribution of alpha activity in the Tactile Only conditions, with more posterior alpha (*p* = 0.047) compared to controls (while groups did not significantly differ in conditions with N-back tasks, *p* = 0.155). Meditators showed an altered distribution of alpha in the Tactile Only conditions compared to N-back conditions (*p* < 0.001, η² = 0.19). Controls also differentiated significantly between conditions, however, this change is smaller than the change observed among meditators (*p* = 0.046, η² = 0.04). B – MDS analysis comparing mean alpha topographical maps between groups and Tactile Only/ N-back conditions. The graph indicates that meditators showed more parietal occipital activation during the Attend Tactile Only and more frontal activation during the conditions with N-back. *Note*: N-Back = average of conditions with the N-back Task, Tactile Only = average of conditions without the N-Back task.

## Discussion

The present study examined whether experienced mindfulness meditators showed differences in neural resource allocation under single and cross-modal task demands. The conditions allowed for examination of differential processing of visual and somatosensory stimuli that required processes including WM, sustained attention, and divided attention. The combination of these processes has not been previously studied in meditators. Meditators showed a P300 with more fronto-central positivity when responding to visual stimuli, and greater ability to modulate alpha distribution between low (tactile-only condition) and high (dual task condition) task demands requiring more neural resources. These differences in neural activity were concurrent with improved WM performance in the meditator group. The results provide evidence of differences in neural activity and resource allocation between meditators compared to demographically-matched controls.

### Improved WM

Existing literature has shown positive effects of mindfulness practice on WM, attention, and cognitive control functions (Mrazek, Franklin, Phillips, Baird, & Schooler, 2013; Quach, Jastrowski Mano, & Alexander, 2016; Zeidan et al., 2010). The present study adds evidence of improvements in WM functions during divided attention by demonstrating that meditators performing with higher accuracy on the N-back task while simultaneous attending to distractor stimuli. The findings indicate an enhanced ability for meditators to inhibit distraction and distribute neural resources during cognitively demanding task conditions.

### More frontally distributed WM ERPs in meditators

Neural distribution differences between groups during the P300 window are consistent with previous work by Bailey et al. (2018), whom identified a similar pattern of frontal P300 distribution among meditators along with enhancements in attentional control and related behavioural performance during the Go/No-go task. Unique to the present study, source analysis (see supplementary materials 1) demonstrated that meditators demonstrated greater neural activity in visuospatial processing regions and less activity in somatic (sense of touch) processing regions in response to visual WM stimuli (Kim et al., 2015; Trés & Brucki, 2016). The capacity-limited nature of WM functions by modulating neural activity to attend to task-relevant processing regions, and use disengagement mechanisms to guard against interference, enhancing task-relevant information processing (Jha et al., 2019; Shipstead et al., 2016; Sreenivasan & Jha, 2007). Findings from this study demonstrate that meditators likely attended to the N-back visual stimuli more strongly in both low and high resource demanding conditions, suggesting enhanced ability to orient neural activity to the visual task while simultaneously disengaging from the tactile stimuli.

The enhanced ability among meditators to orient neural activity in favour of task-relevant processing regions is likely to reflect mindfulness practice effects on WM encoding, attention, and cognitive control functions (Kok, 2001; Polich, 2007). In a series of studies on WM and cognitive control, Lavie, de Fockert, and colleagues (de Fockert and Lavie, 2001; Lavie et al., 2004; Lavie, 2005; Lavie and de Fockert, 2005) demonstrated that distractor processing increased with greater WM load, and WM load predicted selective attention performance. Using a dual-task study design, participants performed a WM task (memorising a string of numbers) with gradual loading from low to high difficulty (more complex strings of numbers) while simultaneously attending to a visual distractor (e.g. distractor faces). The results showed that increased WM load resulted in greater distractor processing and interference on task performance. The authors theorised that distractor perception is reduced in favour of task relevant processing, however, its ability to guard against interference began to deteriorate with increasing WM demand due to diminishing cognitive control. On this account, the differences in neural activity in meditators in the present study suggest greater capacity in managing high WM loads compared to controls, without compromising distractor reducing functions, suggesting that mindfulness practice may result in more efficient allocation of cognitive control functions during resource demanding tasks.

It was also hypothesized that meditators would show enhanced P300 amplitudes in response to visual WM stimuli, as we assumed this would reflect increased neural resource allocation among meditators when processing the WM stimuli. However, contrary to our hypothesis, no group differences were found in global P300 amplitude during WM conditions. Previous studies in mindfulness-meditation focused on local (single electrode or region specific) P300 modulation effects (Delgado-Pastor, Perakakis, Subramanya, Telles, & Vila, 2013; Eddy, Brunyé, Tower-Richardi, Mahoney, & Taylor, 2015). Single electrode analyses present challenges in analysing neural activity as they cannot discriminate between neural activity distribution and strength of neural response (Koenig et al. 2011). If a single frontal electrode was selected for analysis in the current study, the results were likely to have shown increased P300 amplitudes in the meditator group, as those electrodes showed increased positive voltages compared to controls.

P300 amplitudes can vary across electrodes depending on brain region, with frontal and parietal P300s representing unique cognitive processes (Kilner 2013; Kok, 2001; Polich, 2007). Frontal P300s are often related to decision relevant processes (Polich, 2007; Verbaten, Huyben, & Kemner, 1997), whereas parietal P300s generally represent context updating and memory processes (Kok, 2001; Linden, 2005; Polich, 2007). Previous research investigating WM ERPs using the N-back task have consistently found reduction in P300 amplitudes with increasing memory loads at parietal regions (Causse, Fabre, Giraudet, Gonzalez, & Peysakhovich, 2015; Dong, Reder, Yao, Liu, & Chen, 2015; Scharinger, Soutschek, Schubert, & Gerjets, 2015; Scharinger, Soutschek, Schubert, & Gerjets, 2017). Compared to frontal regions, memory load tended to have a greater effect on parietal regions, and this effect was associated with reduced WM performance (Dong et al., 2015). Attention must be divided between updating (retrieval and replacing WM contents) and inhibition (supressing irrelevant sensory information) processes during demanding tasks. Reductions in parietal P300 amplitude represent less resource allocation to updating and inhibition functions (Scharinger et al., 2015) resulting in poorer behavioural performances (Scharinger et al., 2015). However, in situations where parietal activity and functioning declined (for example old age), greater recruitment of resources from frontal regions have been observed to compensate for parietal decline. Frontal compensation is marked by consistent frontal P300 amplitudes and unchanging behavioural performances (van Dinteren et al., 2014). The frontal compensation might indicate that context updating and memory processes during cognitive tasks can be executed by different brain regions resulting in similar behavioural results. Moreover, these findings suggest that WM functions and performance are likely not represented by a single spectrum of neural activity.

Results from the present study might indicate differences in the unique neural processes employed by meditators during cognitively demanding tasks. Meditators in the present study demonstrated greater positive neural activity in frontal regions and less activity in parietal regions during the P300 window, mirroring compensatory neural activity and comparable behavioural performance from studies among the elderly. However, in the current study this pattern of activity was concurrent with improved behavioural performance in the meditator group. In view of this, our results might suggest that meditators were not simply more proficient at modulating typical WM related neural activity, but that they engaged in a qualitatively different neural process. Greater frontal activity suggests that when performing WM tasks, meditators may have engaged more frontal regions to process and execute task demand and relied less on parietal functions to support related cognitive functions.

### Alpha Modulation and Working Memory

Meditators showed more alpha activity in parietal-occipital regions in Tactile Only conditions (non-WM), indicating greater inhibition of non-relevant visual information processing when the task did not require visual information to be processed. This result is in-line with previous work by Kerr et al., (2013), who theorised that meditators were likely to show enhanced anticipatory control over somatotopic alpha rhythms, and thus exhibit greater top-down alpha modulation over sensory brain regions. The current results extend the work by Kerr et al. (2013) to visual processing regions. The proposed alpha modulation is consistent with evidence of enhanced attentional regulation and modulation of somatosensory alpha rhythm after short periods of meditative practice (Jha et al., 2007; Kerr et al., 2011). However, while mediators showed more global alpha activity in the Tactile Only conditions, alpha activity to the tactile stimuli did not increase during conditions with the N-back task requiring participants to ignore the tactile stimuli. The results indicate that both groups showed less alpha when they were concurrently performing the WM task, and that meditators showed a greater difference in alpha activity between the Tactile Only and N-back (WM) conditions. These results suggest that meditators were inhibiting visual processing in the absence of the N-back task, and then releasing the inhibition during the N-back task when visual processing was required.

The lack of alterations to alpha activity over the somatosensory region during the ‘ignore tactile’ conditions contradicted the initial prediction that meditators would show somatosensory alpha modulation based on attentional demands. This finding also contrasted the results found by Kerr et al., (2013) and may be explained by the equipment used to capture neural activity. Although EEG captures accurate data globally, it is less sensitive to localised data in specific regions (Lopes da Silva, 2013). Thus, EEG may not be equipped to detect precise alpha changes specific to a minor region in the somatosensory cortex. This explanation is supported by the lack of main effect of attend to / ignore tactile stimuli condition, suggesting that the condition manipulation did not differentiate the distribution of neural activity. Kerr et al., (2013) on the other hand adopted magnetoencephalography (MEG), which allows for higher accuracy in capturing local data.

### Limitations and Future Direction

Cross-sectional studies face inherent limitations. Self-selection bias in the present study may reflect inherent differences in neurobiological profiles and personality traits that promote uptake or adherence to mindfulness practice (Mascaro, Rilling, Negi, & Raison, 2013). Additionally, due to the absence of an active control group who are involved in attention intense activities such as learning a language or instrument (Garland & Howard, 2009), the observed differences in attention regulation between groups may be due to the fact that meditators are experienced in a non-specific form of attention training rather than meditation-specific effects. The most conspicuous potential limitation however, is the difference in years of education between groups, which has been shown to affect cognitive performance and function (Park, Choi, Choi, Kang, & Lee, 2018). Tests were performed (see supplementary materials 2) to assess the effect of this confound by replicating comparisons after groups were matched for years of education, excluding the most educated meditators. The results were consistent with the initial analysis suggesting that education level did not explain the group differences found in the present study. Finally, mindfulness meditators were recruited based on self-reported practices rather than adherence to an objective, standardised course or set of practices. Efforts were made to screen participants to ensure that their meditation practice contained the defined practice of focused attention, however styles of practice still varied with in participants due to the wide variety of mindfulness meditation techniques. While this lack of standardization limits the ability to draw conclusions about a definitive set of practices, it allows the generalizability of our results to many forms of mindfulness meditation. It is also worth commenting that the task required prolonged focus and sustained attention with instructions building on each other gradually increasing in difficulty. Amongst other factors, mindfulness-meditation has been considered helpful for reducing mental fatigue (Kaplan, 2001), with evidence of reduced cognitive fatigue after mindfulness practice (Johansson, Bjuhr, & Rönnbäck, 2015). Therefore, performance deterioration due to fatigue may have impacted controls more than meditators and may have influenced the results of the study. However, the lack of interaction between group and condition in accuracy across the two N-back conditions suggests fatigue is unlikely to explain our results.

Future research should consider using MEG to capture neural data with higher accuracy for more specifically localised activity. Additionally, further studies are encouraged to determine whether the differential attentional modulation found in the current study is replicable in studies using a randomised double-blinded longitudinal design to explore meditation effects, which could eliminate a portion of the limitations set out above and help establish causality.

### Summary

The present research extended existing literature by using EEG markers of attentional resource allocation (P300 and alpha activity; Wong et al., 2018) to demonstrate that meditators were able to efficiently allocate neural resources and modulate alpha rhythms to facilitate WM task performance while experiencing a tactile distractor. Attention relevant modulation in somatosensory processing of a tactile stimulation may be particularly significant for understanding mindfulness-specific effects on neural pathways involved in sensory processing, perception, and attenuation (Brown & Jones, 2010; Gard et al.,2012; Nakamura et al., 1997; Zeidan et al., 2011). Overall, the study’s results promote a broader understanding of the mechanisms responsible for the positive effects of mindfulness meditation by demonstrating that mindfulness is associated with objective changes in the brain concurrent with improved WM performance.

## Data Availability Statement

Data sharing was not approved by the ethics committee as participants did not provide consent to sharing the data with third party researchers.

## Function and Declaration of Interests

PBF has received equipment for research from MagVenture A/S, Medtronic Ltd, Cervel Neurotech and Brainsway Ltd and funding for research from Neuronetics and Cervel Neurotech. PBF is on the scientific advisory board for Bionomics Ltd. All other authors have no conflicts to report. PBF is supported by a National Health and Medical Research Council of Australia Practitioner Fellowship 659 (6069070).

## Supplementary Materials

**Supplementary Table 1.** Statistics for d’ scores and Reaction Times in behavioural performance.

|  | Condition | Meditators *M*(*SD*) | Control *M*(*SD*) | Statistics |  |
| --- | --- | --- | --- | --- | --- |
| N-Back *d'* | C4: Attend Tactile,<br>Attend N-back | 2.09 (.51) | 1.90 (.49) | Group Main Effects | $F(1,63) = 4.52, p = 0.038, \eta^2 = 0.07$ |
| | C2: Ignore Tactile,<br>Attend N-back | 2.28 (.42) | 2.04 (.43) | Group x Ignore Tactile, Attend N-back/Attend Tactile, Attend N-back | $F(1,63) = 0.28, p = 0.628, \eta^2 = 0.004$ |
| N-Back<br>Reaction Time | C4: Attend Tactile,<br>Attend N-back | 598.68 (117.17) | 575.29<br>(146.85) | Group Main Effects | $F(1,63) = 0.40, p = 0.528, \eta^2 = 0.006$ |
| | C2: Ignore Tactile,<br>Attend N-back | 552.59 (109.17) | 539.90<br>(120.40) | Group x Ignore Tactile, Attend N-back/Attend Tactile, Attend N-back | $F(1,63) = 0.21, p = 0.648, \eta^2 = 0.003$ |
| Tactile<br>Oddball *d'* | C4: Attend Tactile,<br>Attend N-back | 2.75 (.74) | 2.52 (.87) | Group Main Effects | $F(1,63) = 3.15, p = 0.081, \eta^2 = 0.05$ |
| | C3:Attend Tactile<br>Only | 3.04 (.46) | 2.78 (.64) | Group x Attend Tactile Only/Attend Tactile, Attend N-back | $F(1,63) = 0.01, p = 0.942, \eta^2 < 0.001$ |
| Tactile<br>Oddball<br>Reaction Time | Attend Tactile,<br>Attend N-back | 591.56 (118.11) | 582.04<br>(125.79) | Group Main Effects | $F(1,63) = 1.12, p = 0.295, \eta^2 = 0.02$ |
| | Attend Tactile Only | 491.35 (114.03) | 450.84 (81.23) | Group x Attend Tactile Only/Attend Tactile, Attend N-back | $F(1,63) = 1.18, p = 0.282, \eta^2 = 0.02$ |

### Neural Data; Visual Stimuli Locked ERPs

#### TCT

The TCT was conducted to assess consistency of neural activity within each group and condition separately (Koenig et al., 2011). Although results showed topographical consistency within all groups and conditions during the majority of the entire epoch indicating that TANOVA comparisons between conditions and groups are valid during the majority of time periods. There were some variability within the meditator group in both conditions at different time periods; 527ms to 606ms in the attend condition and 524 to 580ms in the ignore condition. Within the control group, brief periods of variability were observed around the 800 to 900ms mark for both conditions (see figure S1).

**Figure S1.**
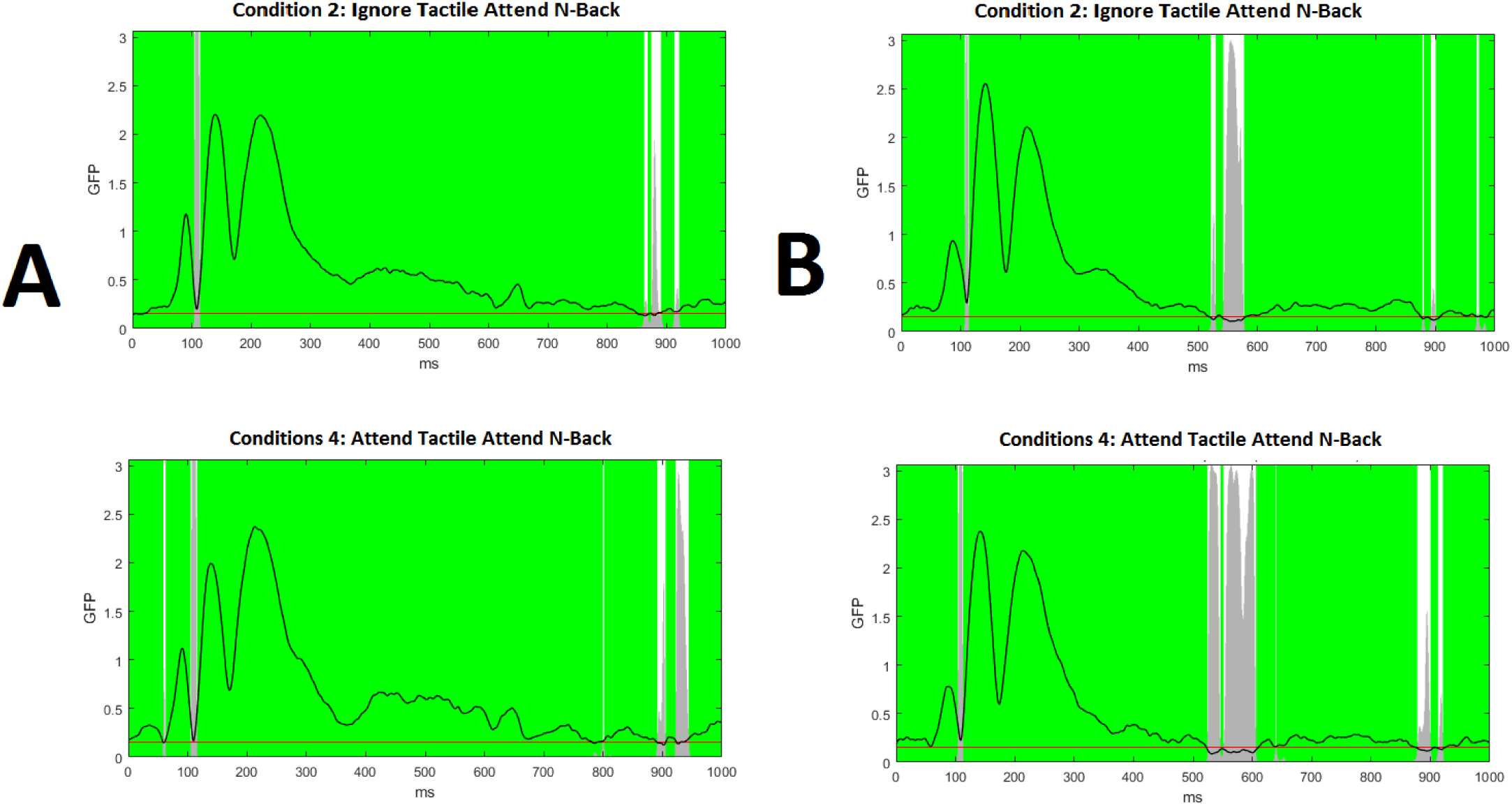
Topographical consistency test. A- Controls. B- Meditators. The GFP values are represented by the line and *p*- values are indicated by grey bars (red line indicating *p* = 0.05). All groups showed topographical consistency across all conditions (marked in green) except for periods after 500ms in both Ignore Tactile, Attend N-back and Attend Tactile, Attend N-back conditions for meditators (marked white), and around the 800 to 900ms mark for the control group in both conditions.

#### Microstates

A microstate analysis approach was used to further explore differences in ERPs by clustering different time periods into prototypical scalp topographies. The interpretation of these results was restricted to durations showing significant group main effects in the TANOVA (from 299 to 352ms and from 419 to 613ms). For group effects, microstate 4 shows significant group main effect for area under the curve (*p* = 0.004, controls 298ms x μV, meditators 128.9ms x μV) and for duration (*p* = 0.004, controls 353ms, meditators 77ms). Meditators showed microstate 4 only once around 200ms post stimulus and then transitioned to microstates 8, 6, and 7 successively. In comparison, controls showed an increased duration of microstate 4 around 200ms post stimulus, then returned to microstate 4 twice around 400ms and 500ms post stimulus. Meditators showed an increased duration of microstate 5 (*p* = 0.014, meditators = 141ms, controls = 45ms), a finding that was driven by the meditator group showing more of this microstate from 540ms onwards, while the control group only showed microstate 5 briefly after the 650ms mark. A significant main effect of group in duration of microstate 8 (*p* = 0.022) was also found, with meditators (164ms) showing greater duration than controls (33ms) (this finding was within the second window of significance in the TANOVA). Figure S2 depicts microstate differences between groups.

**Figure S2.**
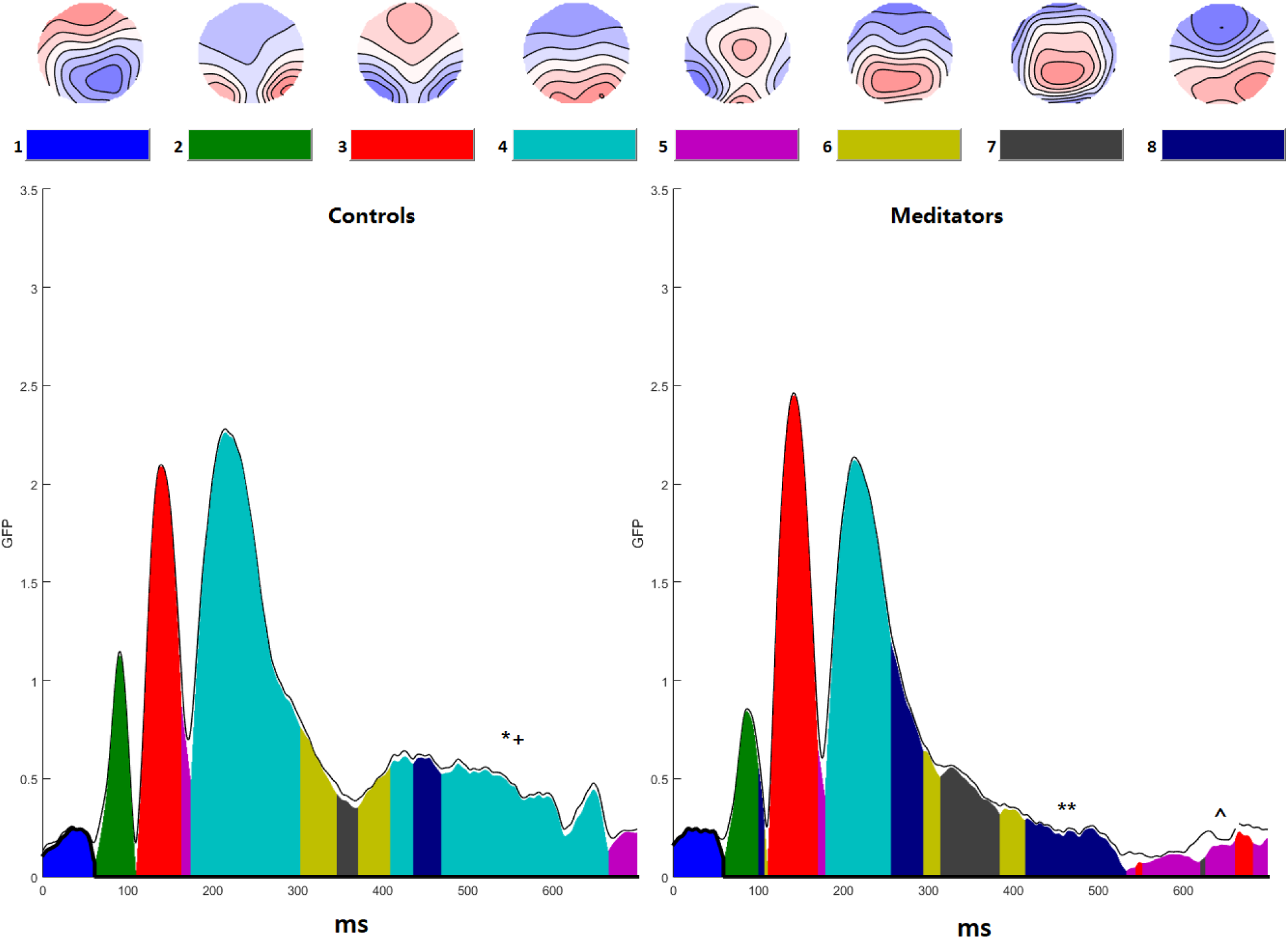
Microstate analysis showing overall between-group effects. Meditators differed in microstate 4, 5, and 8, reflecting first and second significant windows in the TANOVA. * *p* < 0.01 indicates a larger area under the curve in controls. + *p* < 0.01 indicates a longer duration in controls. ^ *p* < 0.05 indicates larger area under the curve in meditators. ** *p* < 0.05 indicates longer duration in meditators.

#### Source Localisation and Pre-processing

Task related cortical sources for ERP data were estimated using Brainstorm (Tadel, Baillet, Mosher, Pantazis, & Leahy, 2011) (http://neuroimage.usc.edu/brainstorm/). Since MRIs were not a part of the study, EEG data were registered with template model ICBM 152. The forward model used OpenMEEG software which adopts the Symmetric Boundary Element Method (Gramfort, Papadopoulo, Olivi, & Clerc, 2010) and the inverse model used Standardized low-resolution brain electromagnetic tomography (sLORETA) to normalise activity (Pascual-Marqui, 2002). The P100 occipital ERP (averaged from 50-150ms) was localised to the correct position to ensure reliability of the source analysis without MRI templates (Malinowski, Moore, Mead, & Gruber, 2017). Topographical analysis at the scalp level showed significant differences in neural activity. Accordingly, source statistical comparisons were not performed (without MRI co-registration source analyses are less reliable) (Michel et al., 2004).

#### Source Analysis

sLORETA was used to estimate cortical sources of the ERP signal time-locked to the N-back stimuli. Source analysis suggested that meditators showed more activity in the left and right posterior parietal cortex during the first (299-352ms) significant window found in TANOVA. This area is thought to be a generator of the P3 (Polich, 2007). Meditators showed less activity during the second significant window found in the TANOVA (419-524ms) in a broad area of the cortex including the right lateral occipital gyrus, supplementary motor area, and SMA of the superior frontal gyrus, as well the posterior section of the left superior temporal gyrus (STG) and middle temporal gyrus. Additionally, meditators showed more activity in the left posterior parietal cortex during this time window. Figures S3 and S4 depicts source reconstructions of the first and second significant window found in TANOVA.

**Figure S3.**
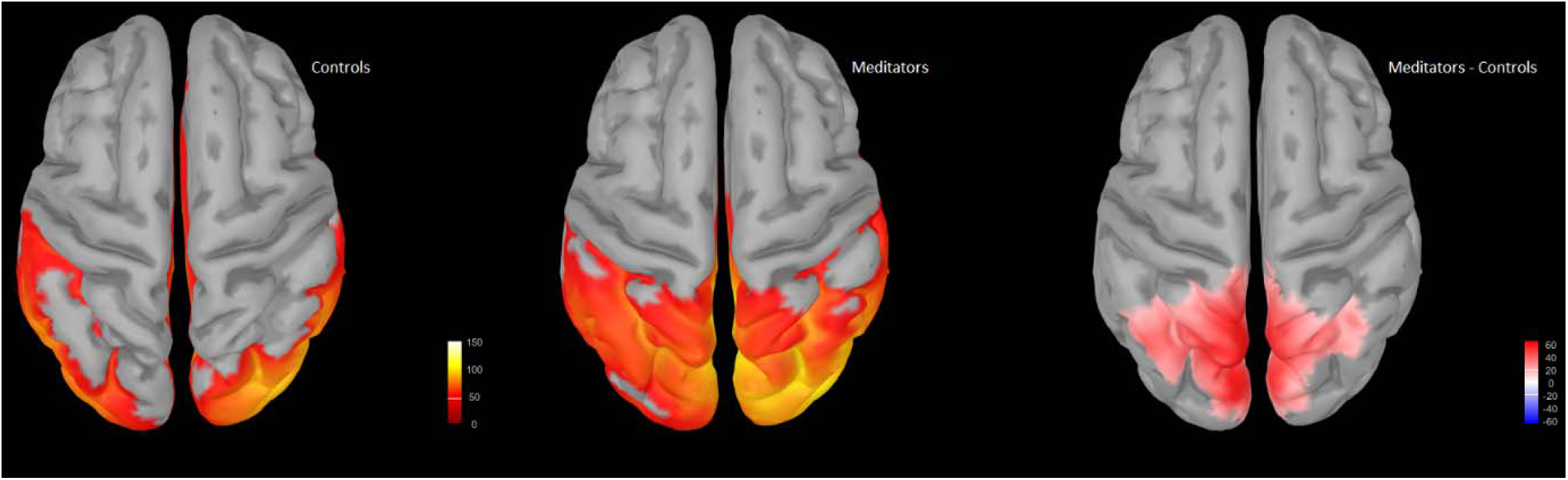
Source reconstruction during the 299 to 352ms window using sLORETA and minimum norm imaging. Please note, only activated brain regions are displayed in group averages and do not reflect positive or negative voltages. Difference maps reflect meditator minus control activity, with red reflecting greater activity in meditators and blue reflecting less activity in meditators, compared to controls.

**Figure S4.**
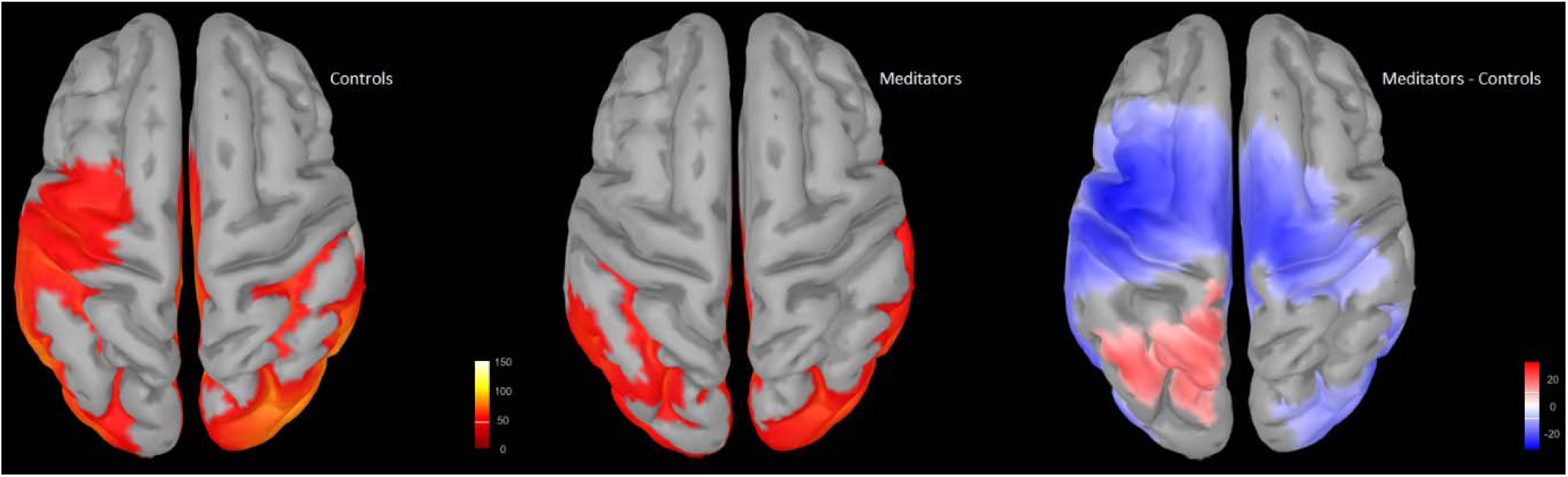
Source reconstruction during the 419-524ms window using sLORETA and minimum norm imaging. Please note, only activated brain regions are displayed in group averages and do not reflect positive or negative voltages. Difference maps reflect meditator minus control activity, with red reflecting greater activity in meditators and blue reflecting less activity in meditators, compared to controls.

## Supplementary Materials 2

Significant comparisons were replicated after excluding the six meditators with the highest years of education so that groups did not differ on years of education. For N-back d’, a significant main effect of group was found (F(1,57) = 5.018, *p* = 0.029, η² = 0.081). A significant effect of group was also found in the TANOVA comparing averaged activity from 299 to 352ms (*p* = 0.01080, η² = 0.064). A significant effect of group was also found in the TANOVA from 419 to 524ms (*p* = 0.0024, η² = 0.086). Figure S5 depicts topographical differences between groups for the first significant window from 299 to 352ms, and for the second significant window from 419 to 524ms. With regards to alpha activity, a significant interaction was found in RMS averaged across the 0 to 1000ms epoch between group and WM task present/absent (*p* = 0.0036 η² = 0.059, ηp² = 0.099, see Figure S6). A significant interaction was found in TANOVA between group and whether participants were completing the N-back memory task or not (*p* = 0.0002, η² = 0.073, η² = 0.080, see Figure S7). Note that the pattern of the significant group main effects for behavioural data and ERPs and interactions for alpha activity was the same as the results from the main comparisons (which included all participants) with stronger effect sizes.

**Figure S5.**
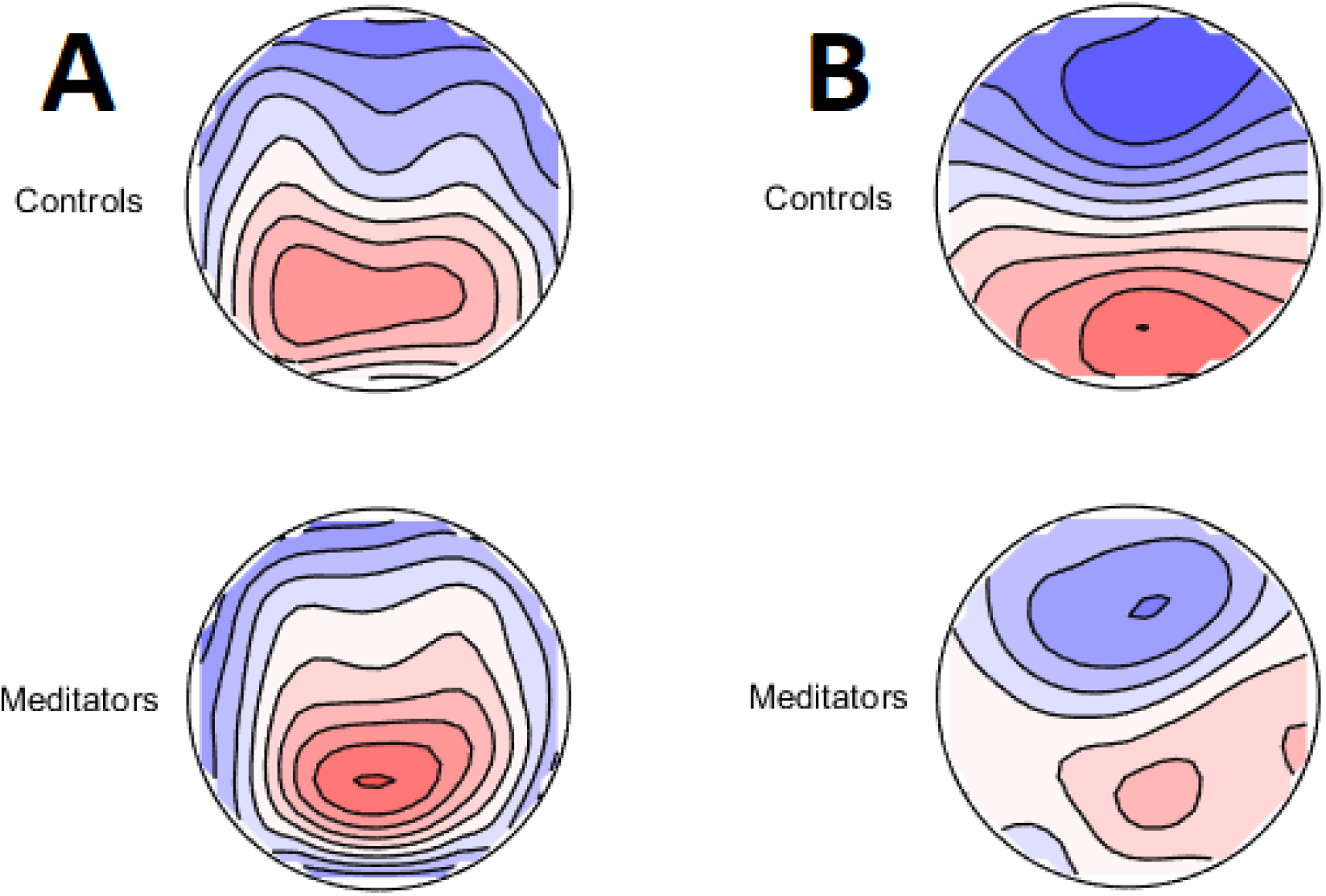
A - Topographical differences between groups for the group main effect in the first significant window from 299 to 352ms (*p* = 0.01080, η²= 0.064). B – Topographical differences between groups for the group main effect in the second significant window from 419 to 524ms (*p* = 0.0024, η²= 0.086).

**Figure S6.**
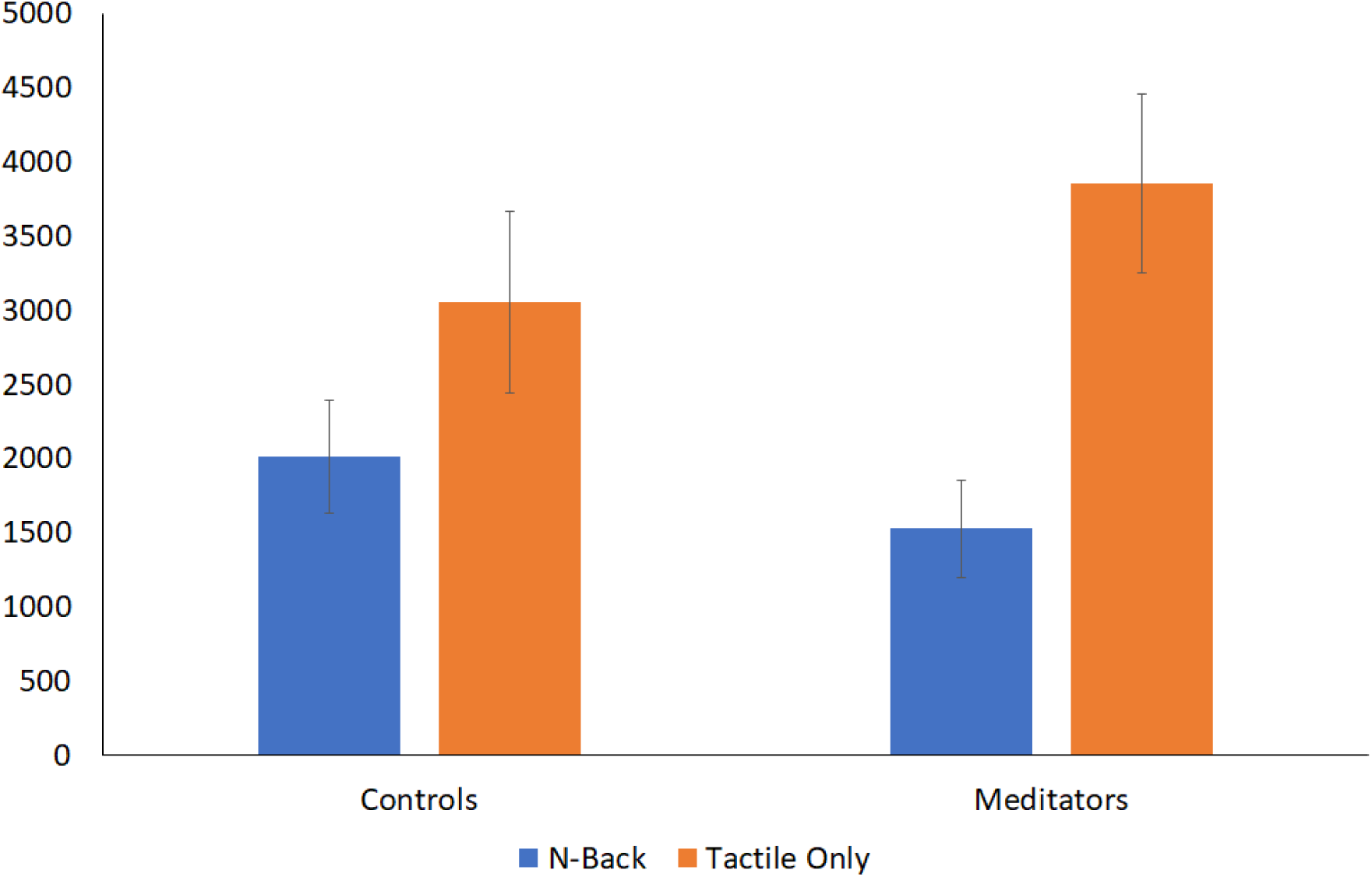
Averaged alpha activity RMS values across the significant time period 0 to 1000ms. Meditators showed a greater difference between conditions than controls (*p* = 0.0036, η² = 0.059, ηp² = 0.099). *Note*: N-Back = average of conditions with the N-back Task, Tactile Only = average of conditions without the N-Back task.

**Figure S7.**
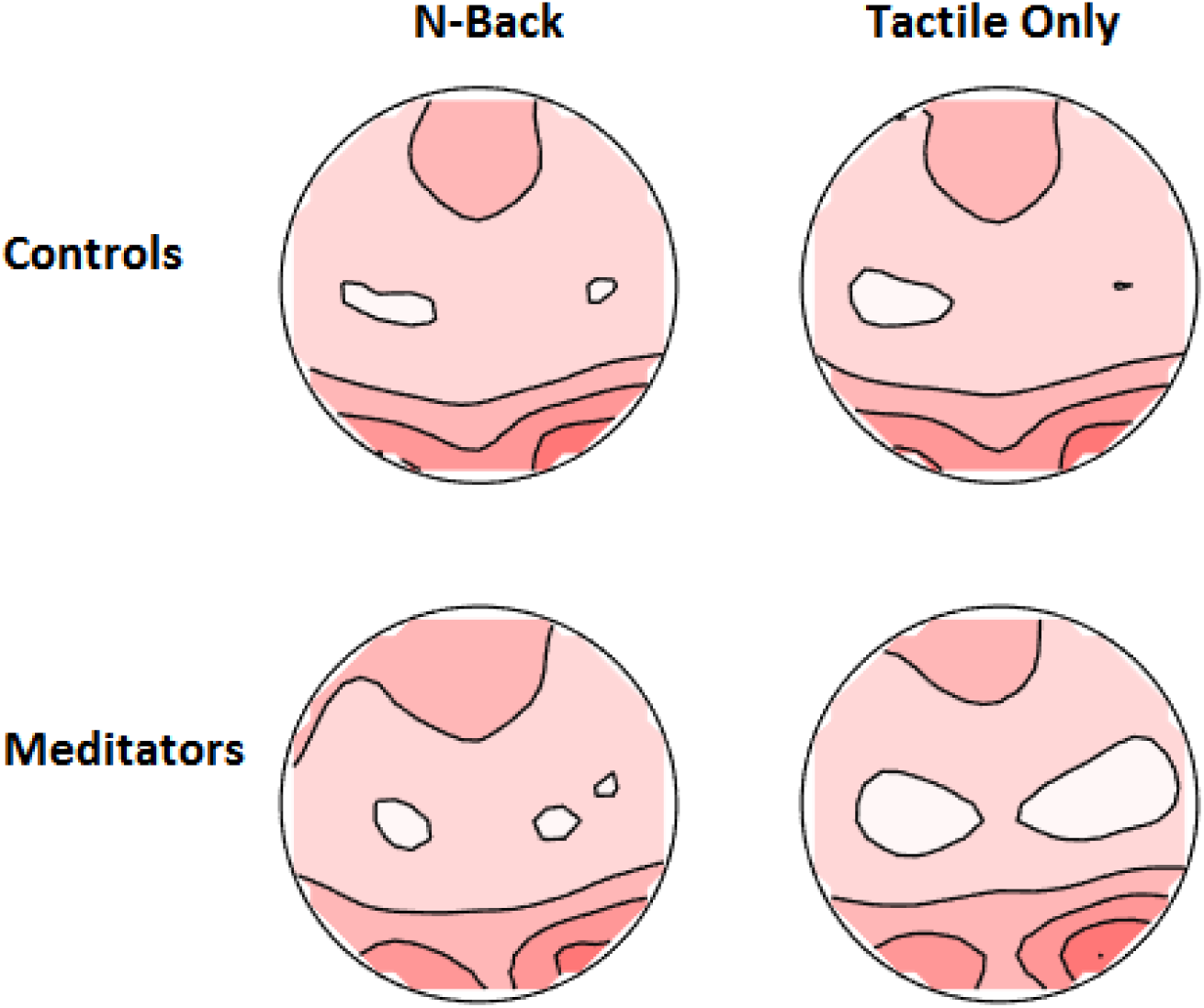
Averaged alpha topographies of Group x N-back/Tactile Only conditions averaged across the 0 to 1000ms window. A significant interaction was found between group and whether participants were completing the N-back memory task or not (*p*-uncorrected = 0.0002, η² = 0.073, ηp² = 0.080). Note that the pattern of the significant group main effects for behavioural data and ERPs and interactions for alpha activity was the same as the results from the main comparisons (which included all participants) with stronger effect sizes. *Note*: N-Back = average of conditions with the N-back Task, Tactile Only = average of conditions without the N-Back task.

